# Revealing genomic changes responsible for cannabinoid receptor loss in parrots: mechanism and functional effects

**DOI:** 10.1101/2022.01.03.474805

**Authors:** Daniel Divín, Mercedes Goméz Samblas, Nithya Kuttiyarthu Veetil, Eleni Voukali, Zuzana Świderska, Tereza Krajzingrová, Martin Tĕšický, Vladimír Beneš, Daniel Elleder, Oldřich Bartoš, Michal Vinkler

## Abstract

In vertebrates, an ancient duplication in the genes for cannabinoid receptors (CNRs) allowed the evolution of specialised endocannabinoid receptors expressed in the brain (CNR1) and the periphery (CNR2). While dominantly conserved throughout vertebrate phylogeny, our comparative genomic analysis suggests that certain taxa may have lost either the CNR1 regulator of neural processes or, more frequently, the CNR2 involved in immune regulation. Focussing on conspicuous *CNR2* pseudogenization in parrots (Psittaciformes), a diversified crown lineage of cognitively-advanced birds, we highlight possible functional effects of such a loss. Parrots appear to have lost the *CNR2* gene at at least two separate occasions due to chromosomal rearrangement. Using gene expression data from the brain and periphery of birds with experimentally-induced sterile inflammation, we compare *CNR* and inflammatory marker (interleukin 1 beta, *IL1B*) expression patterns in *CNR2*-deficient parrots (represented by the budgerigar, *Melopsittacus undulatus* and five other parrot species) with *CNR2*-intact passerines (represented by the zebra finch, *Taeniopygia guttata*). Though no significant changes in *CNR* expression were observed in either parrots or passerines during inflammation of the brain or periphery, we detected a significant up-regulation of *IL1B* expression in the brain after stimulation with lipopolysaccharide (LPS) only in parrots. As our analysis failed to show evidence for selection on altered *CNR1* functionality in parrots, compared to other birds, *CNR1* is unlikely to be involved in compensation for *CNR2* loss in modulation of the neuroimmune interaction. Thus, our results provide evidence for the functional importance of *CNR2* pseudogenization for regulation of neuroinflammation.

## Introduction

Pseudogenization, leading to gene loss, is a common phenomenon in organisms and is responsible for evolutionary changes in immunity and other physiological functions (Wang et al. 2006). Gene loss may be involved in adaptive responses to environmental or pathogen-driven changes in selective pressures (Olson 1999) or represent a random shift in gene content with deleterious effects insufficient to be prevented by negative selection (Charlesworth 2012). Genomic chromosomal rearrangement is likely to be an important source of gene loss events. Massive karyotype alterations have profoundly affected vertebrate evolution in general (Damas et al. 2021), as well as the evolution of certain crown lineages (Nanda et al. 2007; Harewood and Fraser 2014; Furo et al. 2018). Recent advances in genomic research have allowed thorough mapping of gene losses in a number of gene classes, including immune genes such as immune receptors, cytokines and other molecules directly affecting immune signalling (Wang et al. 2006; Temperley et al. 2008; Bainová et al. 2014; van der Loo et al. 2016; Velová et al. 2018). While this type of analysis is still uncommon in neural regulators of immune function, a genomic database search appears to suggest an interesting pattern of recurrent gene loss in the genes of the endocannabinoid system.

Current research shows that immune defence in the periphery is tightly linked with central nervous system function through several bi-directional pathways. In the first, immune responses triggered by peripheral stimulation (e.g. through gut microbiota) translate into brain activity regulation via multiple neural and hormonal connections (Maranduba et al. 2015). Being involved in the gut-brain axis (Vanner et al. 2016; Acharya et al. 2017), numerous activators (e.g. serotonin, cannabinoids, organic acids) regulate both immune and neural cells to eventually act on the *nervus vagus*, which modulate afferent signals that alter brain functions (Silverman and Sternberg 2012). In the second, peripheral immunity modulates central nervous system functioning through cytokine effects on neural and immune cells (microglia or astrocyte) in the brain (Aguilera et al. 2013). Key roles are played by proinflammatory cytokines, such as interleukin 1 beta (IL1B) (Werman et al. 2004; Rider et al. 2011) which is mainly secreted by immune cells in the periphery but also by brain-based microglia. As such, *IL1B* serves as an important marker of both peripheral and neural inflammation (Bird et al. 2002; Ren and Torres 2009).

The endocannabinoid system is involved in the regulation of both the above-mentioned neuro-immune interplay pathways. Connected to the metabolism of cannabinoids, this system consists of cannabinoid receptors (CNRs), their ligands (endocannabinoids) and enzymes synthesising and degrading cannabinoids (Lu and Mackie 2016). CNRs are coupled with the G_i/0_α protein family, which inhibit the adenylate cyclase and voltage-dependent Ca^2+^ channels and activate K^+^ channels (Nestler 2009). Two receptor types are known, i.e. CNR1, which is mainly expressed in cells of the nervous system, and CNR2, which is mainly expressed mainly in immune cells. The *CNR1* and *CNR2* genes are paralogs found in all vertebrates (Elphick 2012). CNR1 is involved in the regulation of emotions, memory, motor activity, feelings, attention, neuropeptide synthesis and, in birds, singing (Soderstrom and Johnson 2000; Soderstrom and Johnson 2003). However, CNR1 also has an important role in the gastrointestinal tract, where it regulates gut motility and secretion, and even, through hypothalamic cores, energetic metabolism (Cristino et al. 2014; Cani et al. 2016; Greenwood-Van Meerveld 2017). In mammals, CNR2 is expressed most in immune organs, such as the tonsils or spleen, where it typically occurs in NK cells, T and B lymphocytes and macrophages, though it is also expressed in microglia in the brain (Galiègue et al. 1995; Carlisle et al. 2002). CNR2 affects immunosuppression and decreases inflammation, pain and the expression of proinflammatory cytokines, which play an important role in contributing to negative feedback regulation (Maresz et al. 2005; Vincent et al. 2016; Krustev et al. 2017). *CNR2* expression has been shown to increase with activation of immune cells related to higher expression of proinflammatory cytokines (Carlisle et al. 2002), while, Maresz et al. (2005) noted that *CNR2* expression in brain-based microglia of mice was up-regulated during neurological inflammation, contributing to suppression of the inflammatory response.

The *CNR2* gene has remained unidentified in the available assemblies of the budgerigar (*Melopsittacus undulatus*) and kakapo (*Strigops habroptila*) genomes (http://www.ensembl.org). Hence, the aim of this study was to confirm whether *CNR2* is absent in parrots. If so, we hypothesise that parrots will show altered neuroimmune regulation due to the loss of *CNR2*. We use genomic and transcriptomic data to map the putative *CNR1* and *CNR2* loss events across vertebrate phylogeny and, subsequently, reconstruct the *CNR2* loss events in parrots. Finally, by comparing parrot and passerine pro-inflammatory cytokine expression patterns in the brain and other tissues over the course of an immune response, we assess the consequences of *CNR2* loss on neuro-immune regulation in parrots.

## Results

### Identification of the CNR genes in parrot genomes

After downloading all presently available tetrapod *CNR* coding DNA sequences (CDSs) from the Ensembl database (for *CNR1* n = 116, for *CNR2* n = 153, one sequence for each species) and using BLAST to detect missing *CNR* orthologues in these species through the NCBI databases (Table S1), the *CNR* sequence data were used to construct a *CNR* phylogenetic tree visualising *CNR1* and *CNR2* presence and absence (Figure 1). We failed to identify the *CNR2* gene in any parrot species, though it was present in all parrot relatives, including falcons (Falconiformes), seriemas (Cariamiformes) and passerines (Passeriformes). According to Ensembl, the *CNR2* gene is located on the 23rd chromosome in the zebra finch (*Taeniopygia guttata*) and chicken (*Gallus gallus*) genomes, being directly adjacent to the *FUCA1* gene (upstream) and the *PNRC2* gene (downstream; Fig. 2). In the budgerigar genome, we found both these genes on chromosome 14; however, there was a ~5.5 Mbp insertion with inverted gene order directly between *FUCA1* and *PNRC2* (Fig. 2). Using BLAST, we identified short gene fragments showing 28% similarity to the barn owl (*Tyto alba*) *CNR2* and 10% similarity to the blue-crowned manakin (*Lepidothrix coronata*) *CNR2*, 6 072 bp downstream of *PNRC2*. Interestingly, in the kakapo genome, different genes were situated downstream of the *PNRC2* gene (on the 15th chromosome; Fig. 2) and no sign of any remaining *CNR2* gene or pseudogene sequence. To confirm the absence of the *CNR2* gene in parrot genomes, we designed conservative *CNR1* and *CNR2*-specific primers and sequenced the budgerigar gDNA-derived PCR amplicons by Sanger sequencing (Table S2). In contrast to the zebra finch, we found no evidence for *CNR2* presence in budgerigar gDNA. Further, the sequences amplified by the *CNR2*-specific primers from parrot gDNA failing to match any of the sequences annotated in the NCBI databases. Finally, our whole transcriptome cDNA sequencing in skin after experimental induction of inflammation with LPS failed to reveal *CNR2* in budgerigars, or in any of the five other parrot species. We take this as conclusive evidence for the absence of functional *CNR2* in parrots.

**Figure 1.**
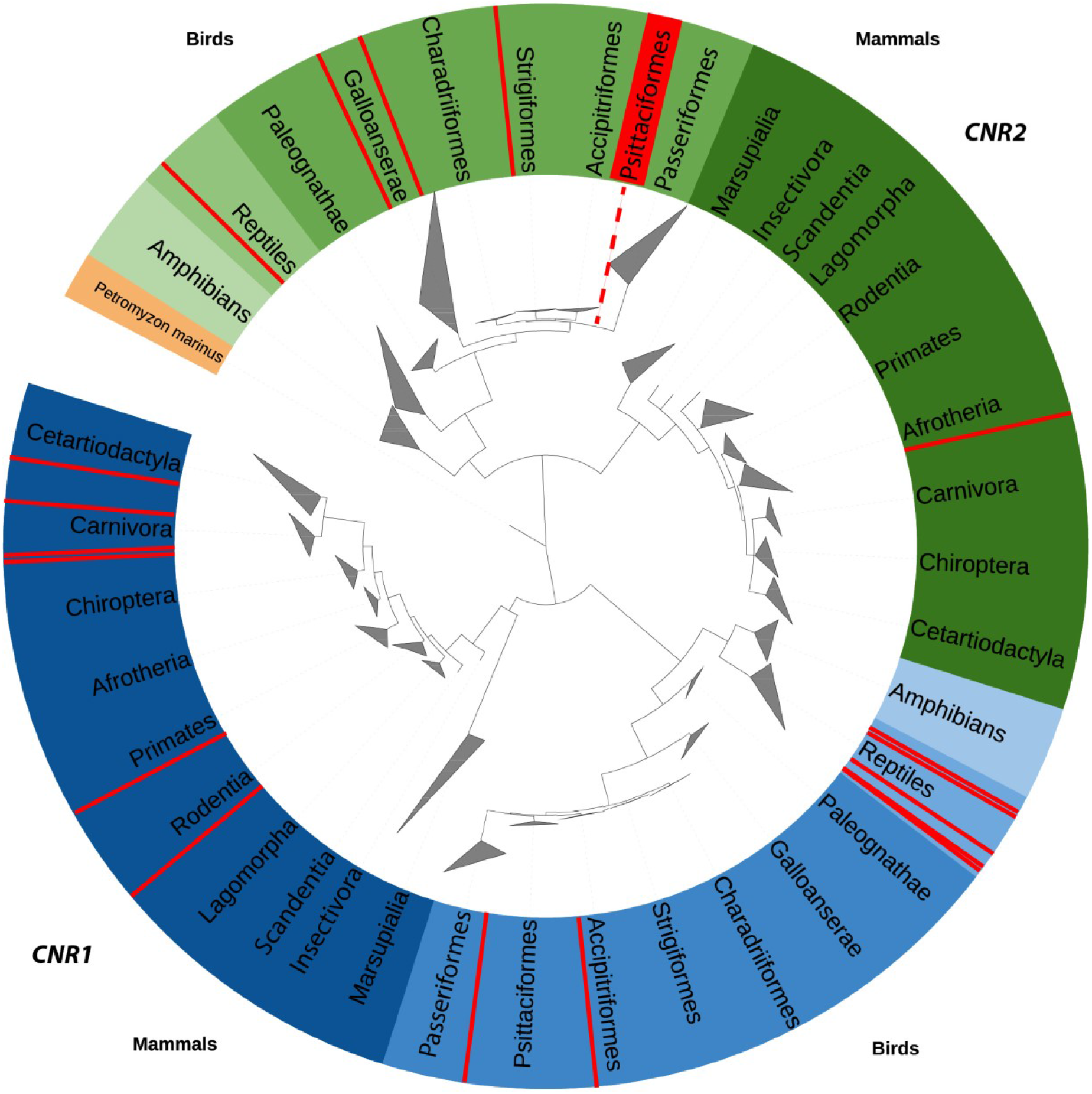
Phylogenetic tree showing gene-specific clustering of *CNR1* (blue) and *CNR2* (green). Lamprey (orange) shows the root of the tree as a common ancestor of the genes. The red colour highlights the presence of species with missing receptors (i.e. cases where the receptors were not revealed by the database search). A fully expanded tree is provided in the Supplemental Material (SM, Figure S1).

**Figure 2.**
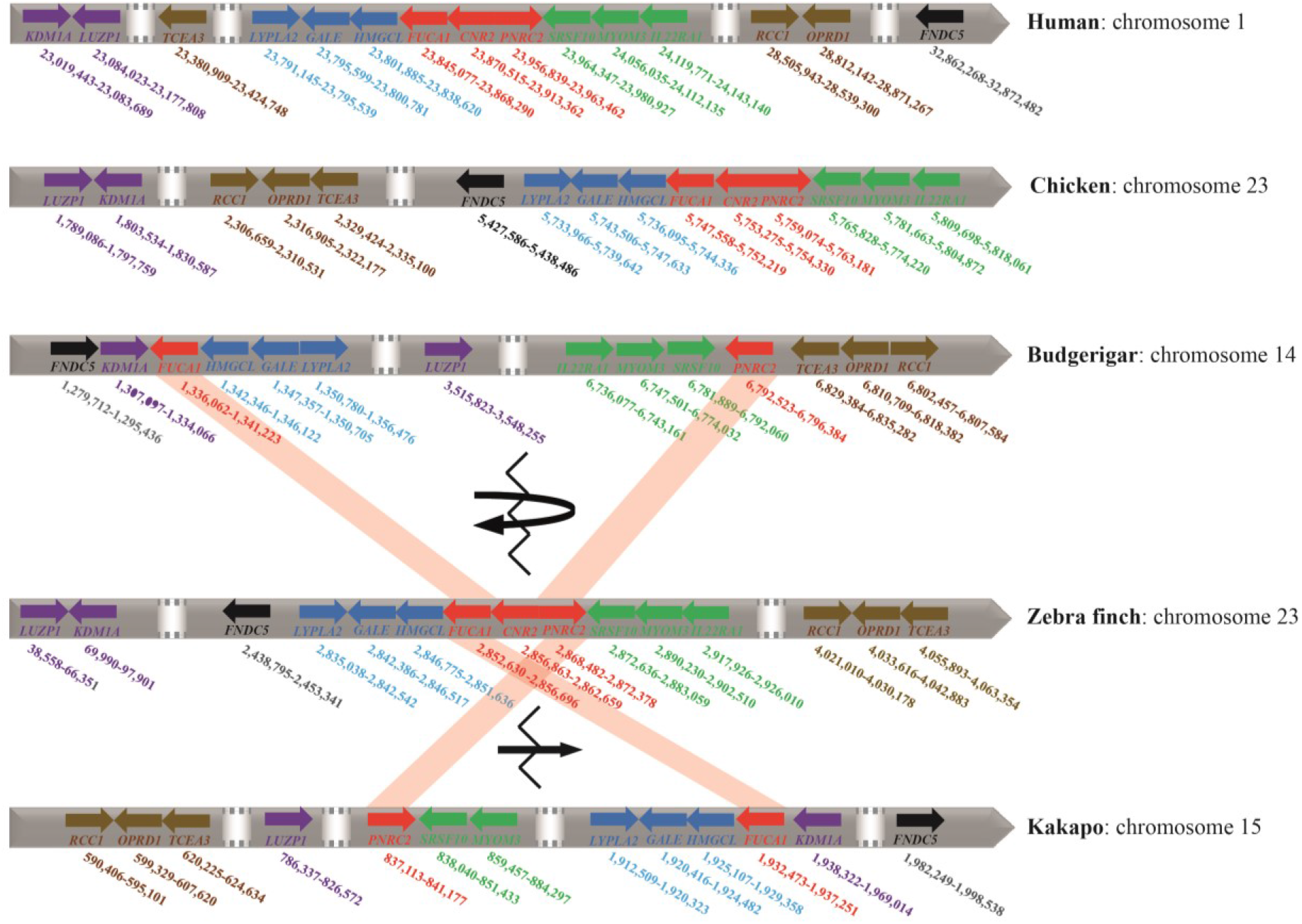
Schematic representation of the *CNR2* locus position and its neighbourhood in the human, chicken, budgerigar, zebra finch and kakapo genomes. *CNR2* and its closest human, chicken and zebra finch neighbouring genes, *FUCA1* and *PNRC2*, are marked in red. The recombination breakpoint in *CNR2* is indicated by a broken line, a curved arrow indicates the inversion event that occurred in the budgerigar evolutionary lineage, while a straight arrow indicates the translocation event that putatively occurred independently in the kakapo evolutionary lineage.

### Positive selection in CNRs

We next questioned the hypothesis that CNR1 took over the functional role of CNR2 in the parrots. Across tetrapods, the test for selection relaxation was not significant in *CNR1* (K = 0.66, p = 0.822, LR = 0.05) and *CNR2* (K = 1.03, p = 0.964, LR < 0.001). Using CONSURF, we identified 67 non-conservative sites in *CNR1* (Fig. 3; Table S5) and 61 non-conservative sites in *CNR2* (Fig. 3; Table S6). The FUBAR (A Fast, Unconstrained Bayesian AppRoximation for Inferring Selection) test failed to identify any positively selected sites in *CNR1*, and only indicated a single site under significant positive selection in *CNR2* (position 7). In comparison, the MEME (Mixed Effects Model of Evolution) test identified seven sites under positive selection in *CNR1* (Table 1); however, changes at these sites only included amino acid alterations in mammals (dogs, bats, squirrels), passerines, amphibians and reptiles, i.e. no specific non-synonymous substitutions with a putatively compensatory role were identified in parrots. In *CNR2*, branch-specific positive selection appeared to be slightly stronger, with MEME identifying 15 sites under positive selection (p<0.10; Table 1). aBSREL (adaptiveBranch-Site RandomEffects Likelihood) also found no evidence of any episodic diversifying selection in parrot phylogeny in the *CNR1* gene or in parrot-related species (i.e. zebra finch, common kestrel) in the *CNR2* gene. PROVEAN (Protein Variation Effect Analyzer), used to identify significant changes in function caused by any amino acid variation, failed to indicate any important changes. Finally, SIFT (Scale Invariant Feature Transform) predicted functional changes in *CNR1* at the sites D466R (with a score of 0.04) and T468I (score 0.04), and in *CNR2* at site V342I (score 0.05); however, none of these changes proved important in any of the avian taxa examined. As such, we consider both CNR1 and CNR2 to be functionally conservative in birds.

**Figure 3:**
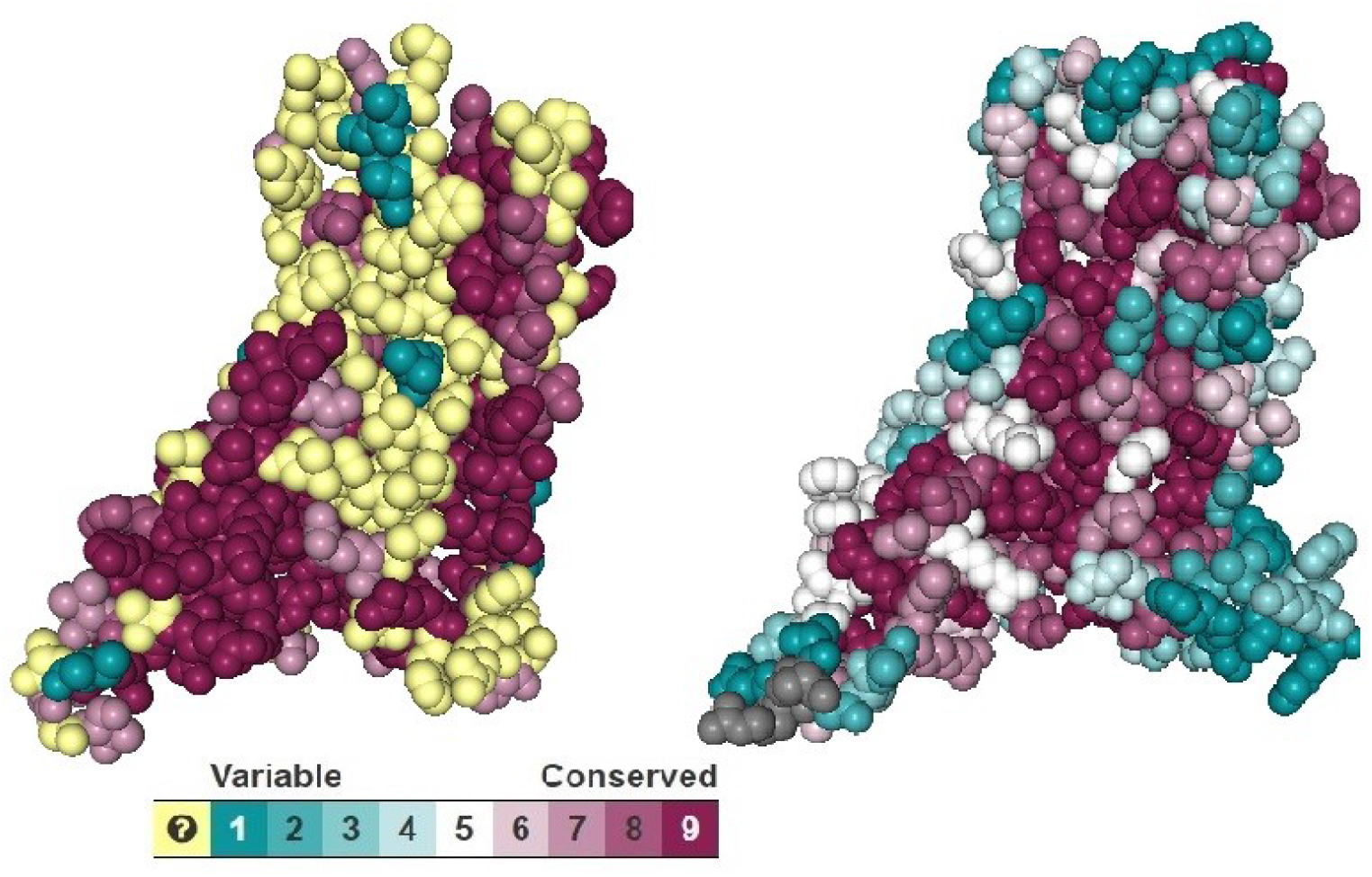
Spacefill models of budgerigar CNR1 (left) and zebra finch CNR2 (right), with transmembrane domains showing variable (blue) and conserved (red) positions. Sites with unresolved conservatism are shown in yellow.

**Table 1:** Amino acid sites identified as under positive selection in *CNR1* and *CNR2* CDS. NS = non-significant; MEME = Mixed Effects Model of Evolution (p-values); FUBAR = Fast, Unconstrained Bayesian AppRoximation (posterior probabilities); Position = position of amino acid in the CNR1 or CNR2 protein sequence

| Sites under positive selection in *CNR1* |  |  |  | Sites under positive selection in *CNR2* |  |  |  |
| --- | --- | --- | --- | --- | --- | --- | --- |
| Position | Substitution | MEME | FUBAR | Position | Substitution | MEME | FUBAR |
| 75 | D/P/E | <<0.001 | NS | 4 | C/E/A/G | 0.01 | NS |
| 463 | V/S | 0.01 | NS | 7 | Y/H/P/F | 0.03 | 0.91 |
| 465 | T/S/G | 0.06 | NS | 19 | G/T/C | 0.01 | NS |
| 466 | D/R | 0.01 | NS | 150 | M/T/V/A/I | 0.07 | NS |
| 478 | T/S/I | 0.01 | NS | 161 | A/T/I/V/F | 0.05 | NS |
| 471 | A/Y | 0.03 | NS | 171 | T/I/V/A/M/K | 0.05 | NS |
| 472 | L/A/V | <<0.001 | NS | 197 | F/L/W | 0.1 | NS |
|  |  |  |  | 208 | I/A/T | 0.07 | NS |
|  |  |  |  | 220 | V/T | 0.02 | NS |
|  |  |  |  | 339 | I/W/V/A | 0.01 | NS |
|  |  |  |  | 342 | V/P/T/I | 0.01 | NS |
|  |  |  |  | 346 | T/V/I/M/D/A | 0.03 | NS |
|  |  |  |  | 347 | R/H/C//T/V/N/<br>I | <<0.001 | NS |
|  |  |  |  | 348 | V/I/C/H/A/T/<br>W/M | <<0.001 | NS |
|  |  |  |  | 349 | M/L/T/V/G/P/<br>D/N/S | 0.04 | NS |

### Transcriptomic evidence for avian patterns of CNR expression during immune response

First, we used previously collected 3’end QuantSeq transcriptomic data from zebra finch skin tissues stimulated *in vivo* with LPS to check for immune-response related gene expression changes in *CNR1* and *CNR2* compared to the inflammation marker *IL1B*. While all three genes could be detected as expressed in the transcriptomic data, we only detected a statistically significant change in gene-specific expression after LPS stimulation (p_ad_ << 0.001) in *IL1B*. In *CNR2*, we observed a marginally non-significant (p_ad_ = 0.055) alteration in expression, while no difference in expression (p_ad_ = 0.998) was observed in *CNR1*.

### Differences in inflammation-associated gene expression changes in parrots and passerines

To assess the functional effects of *CNR2*-loss on immune functioning in parrots, we first followed the *CNR1* and *IL1B* mRNA expression patterns across selected tissues after induction of systemic and local LPS-triggered inflammation in a parrot model species, the budgerigar (see Methods, experiment 1, E1). The resulting tissue-specific and time-dependent dynamics (MAM1, Table 2, Fig. S2), indicated that, compared to controls, relative *IL1B* expression increased in both the brain and other tissues (skin, ileum) following LPS stimulation in treated birds (Fig. 4A, SMAM1-4; p<<0.001, TableS7). In contrast, expression of *CNR1* was independent of LPS treatment (MAM2, Table 2, Fig. S3; Table S7).

**Figure 4.**
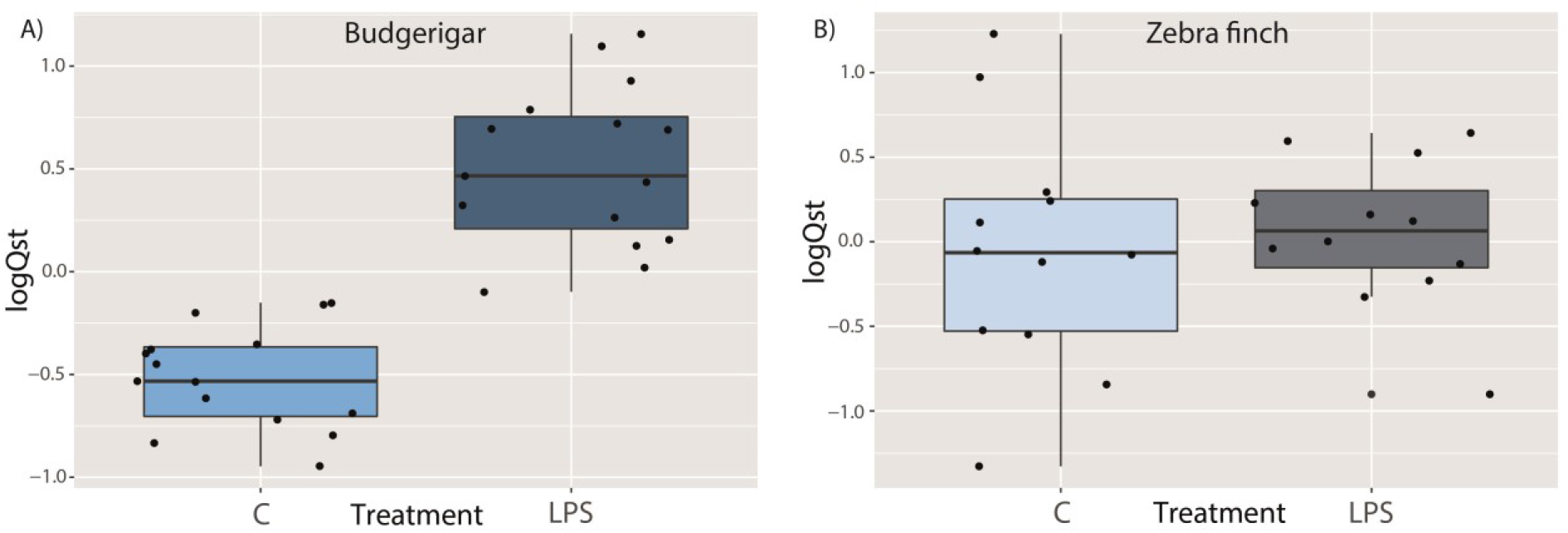
Expression of *IL1B* following peripheral stimulation in the brains of (A) budgerigars and (B) zebra finches. Gene expression is shown as centred standardised relative expression (logQst) values. C = control birds, LPS = stimulated birds

**Table 2:**
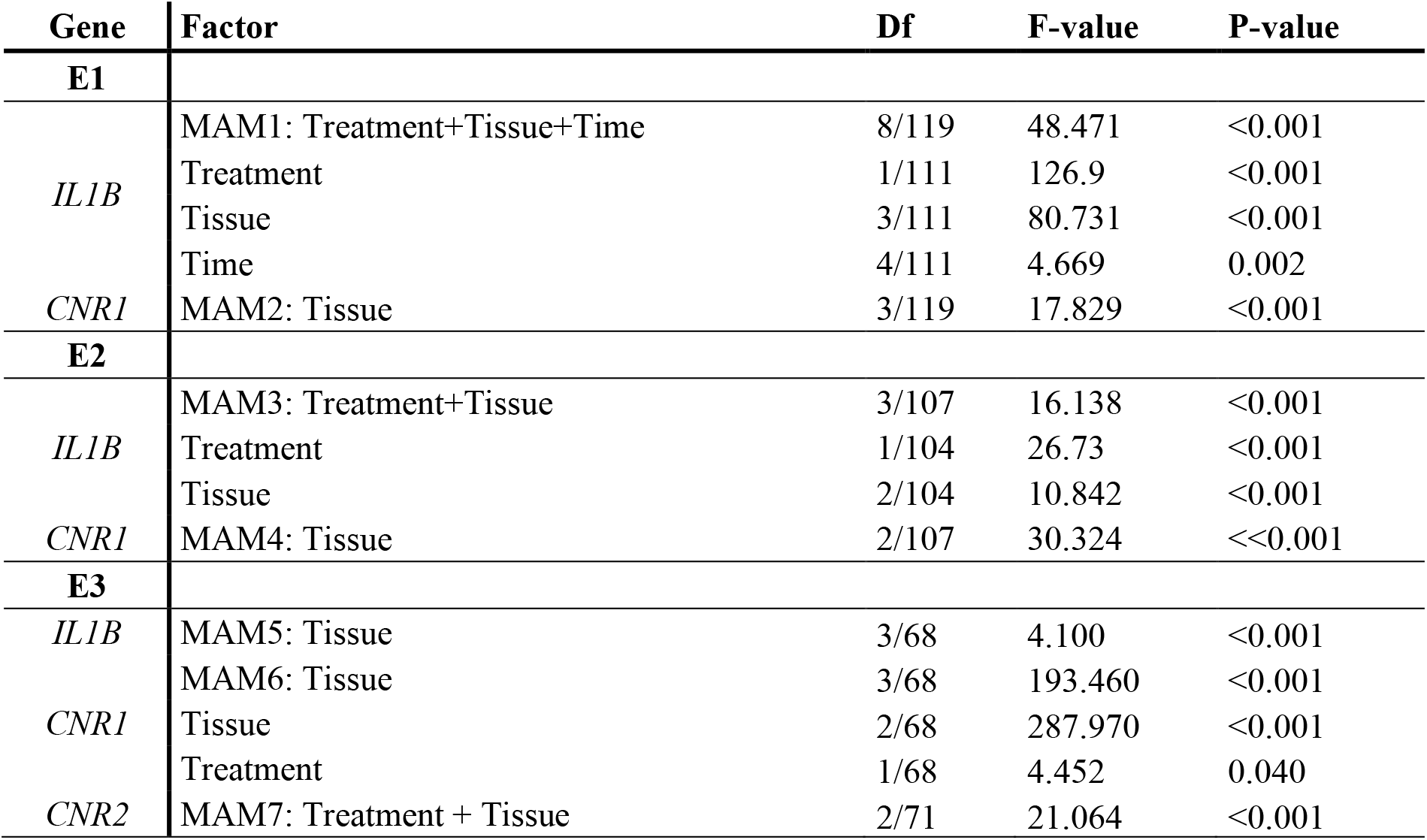
Minimal adequate models for the relative expression (Qst) of selected genes in experiment one (E1), two (E2) and three (E3). MAM = minimal adequate model; DF = degrees of freedom

Given the putative effects of interspecific variability, we next compared changes in *IL1B* and *CNR1* mRNA expression following LPS stimulation in six parrot species (see Methods, E2). The results confirm that expression of *IL1B* depends on LPS stimulation in all parrot tissues, regardless of species (Fig. 5A; MAM3, Table 2; SMAM5-7, Table S8). In comparison, we found no statistically significant effect for LPS stimulation on *CNR1* mRNA expression in any tissue or species (Fig. 5B; MAM4, Table 2; SMAM8-9, Table S8). Furthermore, there was no correlation between *CNR1* expression and expression of *IL1B* (Figs. S4-S6).

**Figure 5.**
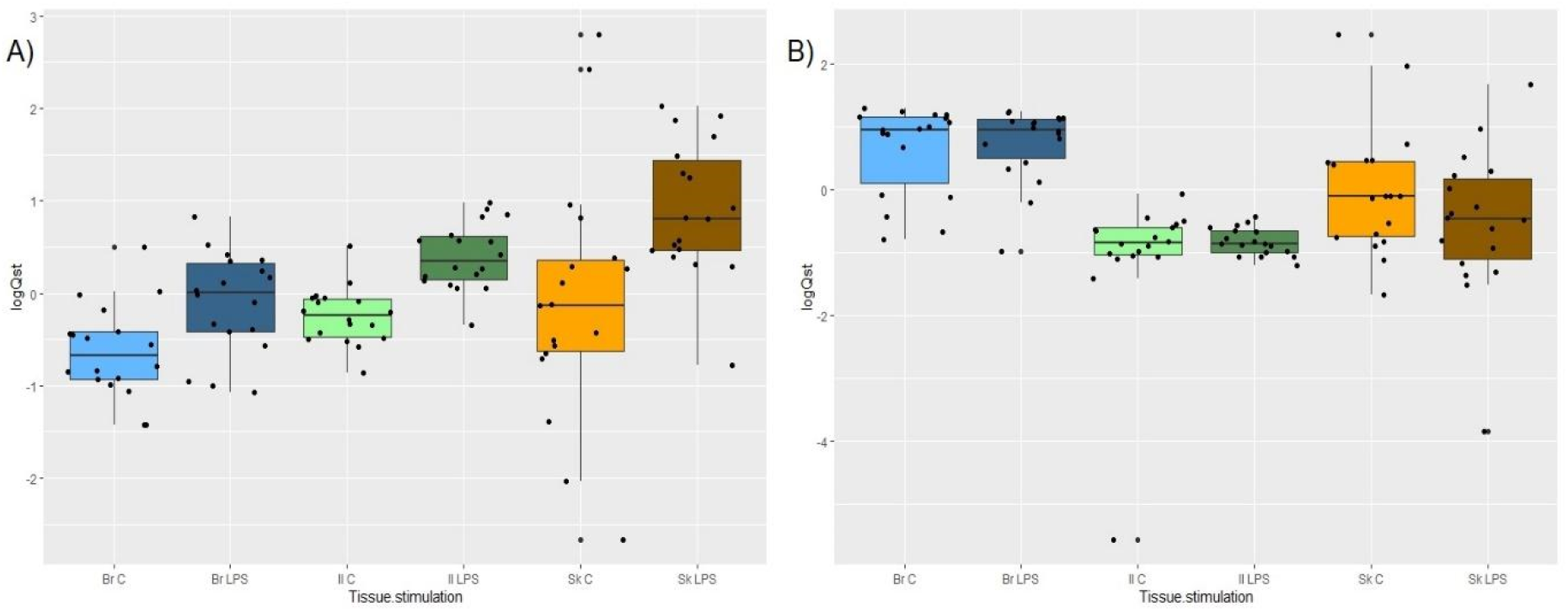
Changes in the level of (A) *IL1B* and (B) *CNR1* expression (Qst values), 24 hours after peripheral stimulation with lipopolysaccharide (LPS), in different parrot tissues. Differences between LPS-treated and control individuals were only statistically significant (p<<0.001) for *IL1B* (not *CNR1)*. Br C = control brain; Br LPS = LPS-stimulated brain; IL C = control gut; IL LPS = LPS-stimulated gut; Sk C = control skin; Sk LPS = LPS-stimulated skin

Finally, to compare brain and peripheral responses between parrots and passerines, we tested for *IL1B* and *CNR1* gene expression changes in the zebra finch, a species known to have a functional *CNR2* receptor. In the zebra finch, *IL1B* expression was also dependent on LPS stimulation in a tissue-specific manner (MAM5, Table 2, Figure S7), with a similar trend also observed in *CNR1* expression (MAM6, Table 2, Figure S8). In contrast, *CNR2* expression was independent of treatment and only differed between tissues (MAM7, Table 2, Figure S9). There was no inflammation-associated change in either *CNR1* (Table S9; Figure S4) or *CNR2* (Table S9; Figure S5) expression in the zebra finch brain. While there was no correlation between *CNR1* and *IL1B* expression in the brain or periphery (Figs. S10-S12), expression of *CNR2* was positively correlated with expression of *IL1B* in the brain (p = 0.009; r = 0.711; Fig. S13), but not in the skin or ileum (Figs. S14-S15). Importantly, we observed no increase in *IL1B* expression in the zebra finch brain following LPS stimulation (p = 0.658, Table S9, Fig. 4B), which is in striking contrast with the up-regulation of proinflammatory *IL1B* in the brain of LPS-stimulated parrots (Figs. 4A and 5A).

## Discussion

Regulation through CNR1 and CNR2 interconnects the signalling of the neural and immune systems. Here, we were able to show that the *CNR2* gene has been lost at least twice during parrot evolution, most probably through chromosomal rearrangement. While absence of the *CNR2* gene has also been reported in other tetrapod species, true loss of the *CNR2* gene requires further confirmation in non-parrot species. Furthermore, we found no evidence for compensatory evolution in *CNR1* after *CNR2* loss in parrots. Interestingly, our comparative experimental findings suggest that these gene loss events probably affect neuroimmune regulation. While mild peripheral inflammation induced by LPS in passerines possessing functional *CNR2* (represented by the zebra finch) failed to trigger any up-regulation in inflammation signalling (measured as *IL1B* expression) in the brain, we recorded significantly increased expression of pro-inflammatory cytokine in parrot (represented by the budgerigar) brains during acute sterile peripheral inflammation.

LPS-induced activation of the immune system in the periphery can trigger systemic immune responses in mammals or birds with neuroinflammatory outcomes (Ban 1992; Sköld-Chiriac et al. 2014; Batista et al. 2019), while in humans and mice, even mild neuroinflammatory changes can cause important alterations in behaviour and cognition (Jurgens et al. 2012). This phenomenon has not been recorded in birds, where even high doses of LPS trigger only mild and non-lethal inflammation (Armour et al. 2020). However, most immunological data for birds has so far only been generated in poultry (representing the evolutionarily basal Galloanserae lineage), or, to a much lesser extent, in passerine birds. Thus, diversity in avian immune responses to peripheral stimulation remains largely unknown. Of particularly relevance to this issue is the investigation of immune response regulation in species with highly rearranged genomes, such as the parrots (Nanda et al. 2007; Furo et al. 2018). Therefore, here we focused on neuroinflammatory effects of LPS-induced peripheral inflammation in budgerigars, a model parrot species, where we revealed *CNR2* pseudogenisation.

Activation of peripheral inflammation can modulate expression of *CNRs* in both the periphery and the brain, thereby altering neuronal processes and behavioural and cognitive functions (Procaccini et al. 2014) during interactions between the nervous and immune systems (Acharya et al. 2017; Vagnerová et al. 2019). We confirmed *CNR1* expression in the nervous system of birds (both zebra finches and parrots), suggesting a similar regulatory effect on neuronal processes as in mammals. However, in this study, we also showed that *CNR1* is also expressed in peripheral tissues, including the ileum and skin. In mammals, leukocyte-modulating *CNR2* is also expressed in both the brain (microglia) and periphery (Maresz et al. 2005); however, somewhat surprisingly, previous radiographic investigations have revealed no signs of its expression in the brain of budgerigars (Alonso-Ferrero et al. 2006). Our genome-database search indicated a complete absence of functional *CNR2* genes in all parrot species investigated, which contrasts with the conservative presence of functional *CNR2* genes in all lineages closely related to parrots (i.e. the falcons, seriemas and passerines, including zebra finches). We were able to identify putative remnants of the *CNR2* pseudogene in the budgerigar genome, however, indicating apparent *CNR2* pseudogenization following massive karyotype rearrangements early in parrot phylogeny (Nanda et al. 2007; Furo et al. 2018). Interestingly, a comparison of the karyotype localisation of passerine *CNR2*-neighbouring genes in the budgerigar and kakapo genomes suggested two presumably independent karyotype rearrangement events in parrots resulting in *CNR2* loss. Absence of *CNR2* was confirmed through negative results for i) *CNR2-*targeted amplification attempts in budgerigar gDNA using conservative PCR primers, and ii) searches through Illumina NextSeq-generated transcriptomes of three tissue types in six different parrot species. We consider this as conclusive support for the complete absence of the *CNR2* gene in parrots.

This finding raises the question as to whether a pseudogenization event could have affected the regulation of neuro-immune interactions in parrots. As we found no other *CNR* gene in parrot genomes aside from *CNR1*, we tested for evolutionary changes in *CNR1* that could be linked to *CNR2* absence. However, our selection analysis showed that this receptor is highly conservative throughout vertebrates and, therefore, no compensatory selection linked to *CNR2* loss is likely in parrot *CNR1*, suggesting *CNR2* pseudogenisation had functional significance. To test for regulatory impacts on parrot immunity from *CNR2* loss, we compared our previously obtained data on systemic inflammation in zebra finches with the newly obtained experimental results for budgerigars. As CNR2 is involved in negative regulation of inflammation-linked *IL1B* expression (Klein et al. 2003; Maresz et al. 2005), we compared both species- and tissue-specific effects of LPS stimulation on the expression of the *IL1B* pro-inflammatory marker. We also measured the expression of *CNR1*, which could be involved in the altered signalling. Similar to other vertebrates (Edelman et al. 2007) and to zebra finches, we observed up-regulation of *IL1B* expression in the peripheral tissues (ileum, skin) after intra-abdominal LPS stimulation in both budgerigars and other parrot species. However, in contrast to the zebra finch, we also observed up-regulation of *IL1B* expression in the brain tissue of parrots when using identical LPS doses. The same pattern of increased proinflammatory signalling (*IL1B* expression) in the brain during peripheral inflammation was detected in both budgerigars and other parrot species, suggesting that parrots in general may be more vulnerable to neuroinflammation than other birds. This is supported by the fact that parrots are exceptionally susceptible to bornavirus-related neuropathy (Rinder et al. 2009; Staeheli et al. 2010; Rubbenstroth et al. 2012; Chen et al. 2020).

CNRs are known to participate in mediating a balance between the immune system and gut microbiota (Acharya et al. 2017) by inhibiting secretion of proinflammatory cytokines and chemokines (e.g. IL8; Hasenoehrl et al. 2016). Our data, therefore, also suggests that *CNR2* loss in parrots impairs the negative regulation of immune responses to symbiotic microbiota, which, in other birds, down-regulates systemic proinflammatory signalling mediated through, for example, IL1B. This corresponds with the fact that *CNR1* expression does not correlate with *IL1B* expression in the brains of parrots or passerines, though there is a correlation in expression of *CNR2* and *IL1B* expression in zebra finches. It would appear that *CNR2* absence results in an increased susceptibility to neuroinflammation, marked by increased expression of *IL1B* in the parrot brain under peripheral stimulation. This pattern is absent in zebra finches, where CNR2 presence may quench systemic inflammation. While our current results must be viewed as an initial analysis of this phenomenon, and thus should still be treated with some caution, current evidence from *CNR2*-knock-out mice showing pronounced immunopathology (Karmaus et al. 2013), appears to support our interpretation. Furthermore, CNR2 has also been shown to provide important anti-neuroinflammatory effects in the human brain (Klegeris et al. 2003; Domenici 2006; Solas et al. 2013; Tao et al. 2016). This not only supports the possible regulatory relevance of *CNR2* absence for sensitivity to neuroinflammation but also suggests that parrots could be prone to dysbiosis-induced neurological syndromes.

## Conclusions

In this study, we provide comprehensive evidence for CNR2 absence in parrots and initial results documenting the possible impact of this loss on regulation of neuroinflammation. Specifically, we observed up-regulated *IL1B* expression in parrot brains, but no similar changes in zebra finches possessing fully functional CNR2. With no apparent compensatory evolution in *CNR1*, parrots lacking functional *CNR2* may be more susceptible to systemic neuroinflammation (e.g. due to dysbeiosis) than other avian species. Our findings not only provide important insights into variability in susceptibility to immunopathology between species but also relevant evolutionary evidence for the functional effects of gene loss events during chromosomal rearrangements.

## Methods

### Identification of CNR-loss events

For the phylogenetic analysis of *CNR1* and *CNR2* genes, we first downloaded all available tetrapod *CNR* CDSs from the Ensembl genome browser database (www.ensembl.org; last accessed on 22.01.2021). Based on a comparison of lists of species with annotated *CNR1* and *CNR2*, we identified all cases of putative *CNR1* or *CNR2* absence. For these species, we performed a targeted search through the NCBI databases (https://www.ncbi.nlm.nih.gov) using blastx and tblastn (https://blast.ncbi.nlm.nih.gov/Blast.cgi) to find the missing orthologues. Using this complete sequence dataset, supplemented with *CNR1* sequences from five other parrots species represented in the E2 obtained by Next Seq Illumina transcriptomic sequencing (see below), we reconstructed the *CNR* phylogenetic tree (based on 318 sequences) in the online tool iTOL to verify the sequence gene-specific orthology (Kumar et al. 2018; Letunic and Bork 2021). (For a list of all species, including their *CNR1* and *CNR2* sequence accession numbers, see Table SI in the supplementary material.) The final dataset consisted of 160 orthologues of budgerigar *CNR1* and 158 orthologues of zebra finch *CNR2* (Table S1). The position of *CNR2* in the zebra finch, chicken and human (*Homo sapiens*) karyotypes was checked in Ensembl and the neighbouring coding genes were identified in parrots with karyotype information available in Ensembl (the budgerigar and kakapo). Based on this data, we reconstructed the genomic changes leading to *CNR2* pseudogenization.

### Selection analysis

We examined evidence for positive selection acting on vertebrate CNRs in order to infer whether loss of *CNR2* was linked to any alteration to *CNR2* functioning over the whole avian phylogenetic clade of parrot-related taxa (including passerines, falcons and seriemas), and whether it resulted in any compensatory evolution in parrot *CNR1*, thereby indicating its functional broadening. First, we used the tool CONSURF (http://consurf.tau.ac.il; (Glaser et al. 2003) to identify non-conservative regions on the CNR surface. Next, we adopted a combination of tools for detecting positive selection available on the Datamonkey server (https://www.datamonkey.org/). All sequences were aligned and truncated to the same length and species with significant insertions or deletions were excluded (41 for *CNR1* and 117 for *CNR2*). Individual sites under diversifying selection were detected using the codon-based maximum likelihood methods FUBAR (Fast, Unconstrained Bayesian Approximation for Inferring Selection; Murrell et al. 2013) and MEME (Mixed Effects Model of Episodic Diversifying Selection; Murrell et al. 2012). Results with posterior probabilities > 0.9 (FUBAR) and p < 0.1 (MEME) were considered significant. To specifically test for episodic diversifying selection occurring in certain phylogenetic lineages, we used the online tool aBSREL (adaptive Branch Site REL; available on the Datamonkey server; Smith et al. 2015) at a threshold of p ≤ 0.05. Finally, the RELAX tool was used to test for relaxation in negative selection on *CNR2* in the parrot-related taxa (Wertheim et al. 2015). We then used the online tools PROVEAN (http://provean.jcvi.org; Choi et al. 2012) and SIFT (https://sift.bii.a-star.edu.sg/; Vaser et al. 2016) to predict functional effects of the amino acid substitutions observed at sites under positive selection.

### Experimental procedures

For the purposes of this study, we used samples and data from two newly performed experiments on parrots and newly analysed data from a previous experiment on zebra finches (Kuttiyarthu et al. in prep.). In experiment 1 (E1), 30 budgerigars were used to map the *CNR1* and *IL1B* expression trajectories of an acute immune response over time. Fifteen individuals were intra-abdominally injected with 0.25 mg of LPS (*Escherichia coli O55:B5;* Sigma-Aldrich, cat. no. L2880) dissolved in 50 μl of Dulbecco’s Phosphate-Buffered Saline (DPBS; Sigma-Aldrich, cat. no. D5652), equivalent to a mean dose of 6 μg of LPS per gram body weight. At the same time, 40 μl of a solution containing 0.2 mg of LPS was subcutaneously injected into the bird’s left wing. For each treated bird, we included one control cage-mate (n = 15). Experimental birds were euthanised with CO2 at different time points, i.e. at 3, 6, 12, 24 and 48 hours post treatment (n = 3 per time point and treatment), and selected tissues were then collected. Blood samples were collected from all individuals and used for subsequent haematological analysis (Bauerová et al. 2020). Haematological analysis was also used to check for health-related effects of manipulation. In experiment 2 (E2), we compared the responses of 36 individuals representing six different parrot species, i.e. the red-rumped parrot (*Psephotus haematonotus)*, the rosy-faced lovebird (*Agapornis roseicollis*), the elegant parrot (*Neophema elegans*), the budgerigar, the cockatiel (*Nymphicus hollandicus*) and the pacific parrotlet (*Forpus coelestis*). The experiment was conducted in three batches with six treatment birds and six controls housed in species-specific pairs per batch. Doses were adjusted to species-specific size as follows: 0.25 mg of LPS dissolved in 50 μl of DPBS per individual for a species of 20–40 g average weight; 0.5 mg of LPS in 100 μl of DPBS per individual of a species of 40–60 g average weight; and 1.0 mg of LPS in 200 μl of DPBS per individual of a species of 80–100 g average weight, each dose being injected intra-abdominally. The left wing web of each treated bird was also subcutaneously inoculated with the following: a dose containing 0.10 mg of LPS in 20 μl of DPBS for a species of 20–40 g average weight; 0.2 mg of LPS in 40 μl of DPBS for a species of 40-60 g average weight; and 0.3 mg of LPS in 60 μl of DPBS for a species of 80–100 g average weight. All birds were euthanised with CO2 24-hours after stimulation and selected tissues collected. Experiment 3 (E3) was based on previously collected data for 24 male zebra finches stimulated using the same methods used in E1 and E2. Briefly, 12 individuals were injected intra-abdominally with 0.1 mg of LPS dissolved in 100 μl of DPBS (6 μg of LPS per gram body weight) and subcutaneously injected into the left wing web with 0.1 mg of LPS dissolved in 20 μl of DPBS. All birds were euthanised by rapid decapitation 24-hours after stimulation and selected tissues collected. All birds from all experiments (E1–3) were obtained from local hobby breeders and housed in pairs in cages 100 × 50 × 50 cm and were left for a minimum acclimation period of three days. The birds had access to food and water *ad libitum* and were kept under a 12L: 12D controlled light/dark cycle with a regulated temperature of 22°C ± 2°C. All birds in all experiments were weighed and divided into two groups (treatment and control) prior to the initiation of experimental work. All tissue samples (skin from the left and right wing-web, ileum and hyperpallial area of the brain) were collected as necropsies after euthanasia and immediately placed into RNA later buffer (Cat. No. 76106, Qiagen, Hilden, Germany), after which they were kept overnight at +4°C and thereafter stored at −80°C until RNA extraction. The research was approved by the Ethical Committee of Charles University, Faculty of Science (permits 13882/2011-30 and MSMT-30397/2019-5) and was carried out in accordance with the current laws of the Czech Republic and European Union.

### Targeted sequencing of IL1B, CNR1 and CNR2 from parrot and zebra finch gDNA

Avian interspecific alignment (Supplemental_Alignment_S1-3) was used to design primers in conservative regions for amplification of the partial coding regions of *IL1B, CNR1* and *CNR2*. PCR amplification was performed using genomic DNA (gDNA) extracted from blood samples (12 samples representing different parrot species, 10 budgerigar samples and 12 zebra finch samples) using the DNeasy® Blood & Tissue Kit (Cat. No. 69581, Qiagen). For PCR, we used the Multiplex PCR Plus kit (Cat. No. 206152, Qiagen), following the manufacturer’s instructions based on a 0.2 μM final primer concentration. gDNA concentration and quality was measured on a NanoDrop 1000 Spectrophotometer (Thermo Fisher Scientific, Waltham, Massachusetts, USA). Using the same set of primers, Sanger sequencing was then performed in the amplicons to assess interspecific and intraspecific genetic variability and to design real-time quantitative PCR (RT-qPCR) primers spanning exon–exon borders and avoiding any inter- or intra-specifically variable sites. To verify assay specificity, cDNAs were amplified using the newly-designed RT-qPCR primer pairs (Table S2). Sanger sequencing was performed at BIOCEV (Biotechnology and Biomedicine Centre in Vestec). The sequences were then quality trimmed, aligned and analysed using Geneious software (http://www.geneious.com, Kearse et al. 2012).

### Transcriptomic search for CNR1 and CNR2 genes in parrots

Small intestine transcriptomes for the six parrot species were obtained from sequencing libraries prepared using the NEBNext Ultra II Directional RNA Library Prep Kit for Illumina (Cat. No. E7760, San Diego, California, USA) in the European Molecular Biology Laboratory (EMBL), Heidelberg (NCBI accession numbers: SAMN23963146, SAMN23963147, SAMN23963148, SAMN23963149, SAMN23963150, SAMN23963151). Paired-end sequencing (80 bp from each end) was then performed on the NextSeq 500 system (Illumina, San Diego, CA) at a sequencing depth of 13–19 million reads per library. Forward and reverse reads were merged, and low-quality reads and adaptor sequences discarded, using BBsuite (‘BBMap’ n.d.). De-novo transcriptome assembly was performed by Trinity (Grabherr et al. 2011) under default settings. To obtain sets of non-redundant transcripts, we applied two filtering steps. First, we used TransDecoder (Haas and Papanicolau 2020) to identify the longest open reading frame of each transcript for each species individually, and second, redundancy was further reduced in the remaining transcript sets by clustering highly similar sequences with CD-Hit (Fu et al. 2012), using a sequence identity threshold of 0.9. Completeness of the six assembled transcript sets against a set of highly conserved single-copy orthologs was assessed using BUSCO (Benchmarking Universal Single-Copy Orthologs v. 4.1.4; Simão et al. 2015). To identify *CRN1* and *CRN2* coding sequences for each species, reference budgerigar (for *CRN1*, Ensembl transcript ID: ENSMUNT00000010298.1) and zebra finch (for *CRN2*, Ensembl transcript ID: ENSTGUG00000001188) sequences were searched using Blastn (Zhang et al. 2000) and compared against raw reads and the sequences obtained for positive selection analysis, and also against transcriptome assemblies.

### Transcriptomic gene expression analysis on zebra finch skin tissue

As an initial check for *IL1B, CNR1* and *CNR2* gene expression changes in zebra finch peripheral tissues, we used the transcriptomic dataset previously sequenced from E3 as part of an unrelated project (Kuttiyarthu et al. in prep.; Accession number: PRJNA751848). Briefly, we used the QuantSeq approach to sequence the 24 male zebra finch skin necropsies, based on 3’end sequencing (Moll et al. 2014), the sequencing taking place at EMBL, Heidelberg. This approach uses whole RNA as the starting material, with no prior poly(A) enrichment or rRNA depletion, the transcripts being generated near to the polyadenylated 3’ end. Samples were first barcoded with Illumina TruSeq adapters and sequencing was undertaken on the Illumina Hiseq 2500 platform. The sequenced samples were then analysed using the BAQCOM pipeline (https://github.com/hanielcedraz/BAQCOM), the adapters being removed using the Trimmomatic tool (http://www.usadellab.org/cms/?page=trimmomatic). The samples were then aligned to the zebra finch reference genome (downloaded from ensemble) using STAR aligner (https://github.com/alexdobin/STAR), the featureCounts (https://www.rdocumentation.org/packages/Rsubread/versions/1.22.2/topics/featureCounts) from the Subread package being used to assign read counts to the genes. Differential gene expression between LPS stimulated and control individuals was calculated using the Deseq2 package in Rstudio v 1.3.1093 (R Studio Team 2021).

### RT-qPCR gene expression analysis

Total RNA was extracted from parrot and zebra finch brain, ileum and skin samples using the High Pure RNA Tissue Kit (Cat. No. 12033674001; Roche, Rotkreuz, Switzerland), the concentration and quality of the RNA being measured on aa NanoDrop 1000 Spectrophotometer (Thermo Fisher Scientific). The RNA was diluted in carrier tRNA (Qiagen, Cat. No. 1068337) enriched with molecular water 1:5 for target genes or 1:500 for 28S rRNA. To calculate the efficiency of each primer pair, a calibration curve was constructed with synthetic DNA standard (gBlocks; IDT, Coralville, Iowa, USA; Table S3) using a dilution series of 10^8^-10^2^ copies/μL, estimated according to Vinkler et al. (2018). The RNA samples and standards were amplified using the Luna® Universal Probe One-Step RT-PCR Kit (E3006, BioLabs®Inc, Ipswich, Massachusetts, USA), with 0.6 mM primer and 0.2 mM probe concentrations (Table S4). RT-qPCR quantification was conducted using a LightCycler 480 PCR platform (Roche) set with the following cycling conditions: (1) 50°C for 10 min, (2) 95°C for 1 min and (3) (95°C for 10 s, 60°C for 30 s) × 45. All assays were performed with template-free negative controls and block positive controls in a freshly prepared dilution series, using *28S rRNA* as a reference gene.

Relative quantification (R) was calculated from the crossing point (Cp) values determined by the second derivative maximum according to Pfaffl (2001), using E and Cp data calculated using LightCycler480 Software v. 1.5.1. To test for gene expression changes between treatment and control birds, we quantified relative gene expression as standardised relative quantities (Qst) according to Vinkler et al. (2018). The PCR efficiencies (E) for parrots were 1.940 for *CNR1*, 1.980 for *28S* and 1.950 for *IL1B;* and 1.967 for *CNR1*, 1.985 for *CNR2*, 2.024 for *28S* and 1.969 for *IL1B* in zebra finches.

### Statistical analysis

All statistical analyses were undertaken using R software v. 3.6.2 (R Core Team 2019). The *Shapiro-Wilk test* was applied to test for data Gaussian distribution. Given their non-Gaussian distribution, Qst and R values were normalised using decadic logarithms. The effects of experimental treatment on gene expression changes was assessed using the ‘Ime4 package’ with linear models (LMs), where gene expression served as a response variable. For the E1 dataset, the full model contained treatment, sex, tissue and time as explanatory variables. As there were important differences between the tissue samples used, we then analysed gene expression changes in the same dataset for each tissue separately using an expanded set of explanatory variables, i.e. treatment, time, sex, tarsus length, initial weight, weight difference (before and after stimulation) and absolute lymphocyte and heterophil counts. For the E2 dataset, the full model contained treatment, sex, batch and species as explanatory variables. For the E3 dataset, tissue and treatment were used as explanatory variables for construction of the full model. Minimum adequate models (MAMs; here defined as models with all terms significant at p ≤ 0.05) were selected by backward elimination of non-significant terms from the full models. All backward elimination steps in the models were verified by changes in deviance with an accompanying change in degrees of freedom (ANOVA) and Akaike information criterion (AIC), using F-statistics. The *correlation test* was used to assess the relationship between expression of *CNR* genes and *IL1B*.

## Data access

All raw and processed sequencing data generated in this study have been submitted to the NCBI BioSample (https://www.ncbi.nlm.nih.gov/biosample/) under accession numbers SAMN23963146, SAMN23963147, SAMN23963148, SAMN23963149, SAMN23963150, SAMN23963151.

## Acknowledgments

We are grateful to Balraj Melepat and Tao Li for their help in the laboratory, and to Kevin Roche for language correction. This study was supported by Grant Schemes at Charles University (Grants Nos. GAUK 646119, PRIMUS/17/SCI/12, and START/SCI/113 with reg. no. CZ.02.2.69/0.0/0.0/19_073/0016935), the Czech Science Foundation (Grant No. P502/19-20152Y) and Institutional Research Support (No. 260571/2021).

## Author contributions

DD and MV conceived and designed the study and drafted the manuscript; DD, MGS, NKV, EV, ZS, TK, MT and MV conducted the experimental work; MGS, NKV and VB performed the sequencing; NKV, OB and MT carried out the transcriptomic analysis; DD and MGS performed the RT-qPCR analysis, and DD carried out the genomic and selection analysis. DE and MV consulted on the interpretation of results. All authors read and approved the final manuscript.

